# Microglia are not required for normal postnatal brain development and cognitive function in the rat

**DOI:** 10.64898/2025.12.22.694658

**Authors:** Emma K. Green, Xinyi Li, Yang Guo, Dylan Carter-Cusack, Rocio Rojo, Maia Lyall, Annie Bell, Helen Zheng, Oliver Teenan, Sally M. Till, David A. Hume, Peter Kind, Clare Pridans

## Abstract

The development, maintenance and survival of microglia, the brain’s resident immune cells, requires signalling via the colony stimulating factor 1 receptor (CSF1R). In mice, the *Fms* intronic regulatory element (FIRE), a highly conserved super enhancer, is required for CSF1R expression in microglia and germ-line deletion of FIRE leads to microglial deficiency. We have generated *Csf1r*^ΔFIRE/ΔFIRE^ (*Fireko*) rats using CRISPR/Cas9. *Fireko* rats are born at expected Mendelian frequency, grow normally and survive to adulthood. Like *Fireko* mice, they lack microglia and subpopulations of resident macrophages in kidney and heart, but retain the other brain associated macrophages and resident populations in the liver, spleen and lungs. In contrast to mutant mice, macrophages in the peritoneum are unaffected and CSF1R expression is not completely abolished in the blood. Aside from the loss of microglia-specific transcripts, the *Fireko* has no effect on expression of genes associated with neurons, oligodendrocytes or astrocytes in adult rats but is associated with detectable calcification in the thalamus. Despite the absence of microglia, we detected no significant differences in ethologically relevant behaviours compared to littermates. The results suggest that microglia are not essential for normal cognitive development. *Fireko* rats provide an alternative to mice as a model for human microglial deficiency and the functions of microglia in development and homeostasis in the brain.

## Introduction

Microglia are the tissue-resident macrophages of the brain. During embryonic and postnatal development and throughout life, microglia are active participants in the removal of superfluous cells and cell fragments including synapses. This process is widely believed to be essential for establishment of neuronal connectivity and activity-dependent synaptic refinement^1–5^.

The proliferation, differentiation and survival of microglia depends upon signals from the colony-stimulating factor 1 receptor (CSF1R) and the two ligands, CSF1 and IL34 (reviewed in ^6,7^). Germ line mutation of the mouse *Csf1r* gene (*Csf1rko*) on a complex genetic background led to the loss of many tissue macrophage populations, osteopetrosis and reduced somatic growth^8^. There have been few subsequent studies of *Csf1rko* mice because of perinatal lethality and hydrocephalus on inbred backgrounds^9,10^. These adverse impacts can be overcome in hybrid F2 mice derived by crossing to an alternative inbred genetic background^11^. Homozygous CSF1R mutations have been generated in rats^12^ and zebrafish^13^ and recognised in a rare human genetic condition^14,15^, in all species leading to complete microglial deficiency.

Heterozygous mutations in the human *CSF1R* gene are linked to a dominant adult-onset leukoencephalopathy^16^. There have now been >200 mutations identified in patients with CSF1R-related leukoencephalopathy^17^. Analysis of mice^18,19^ zebrafish^20^ and patients^21^ supports a dominant negative mechanism whereby mutant receptors interfere with CSF1-dependent activation of wild-type receptors leading to reduced microglial density. The loss of homeostatic microglia in patients with CSF1R-related leukoencephalopathy^22^ is thought to be the predominant cause of behavioural changes and neurodegeneration^17^.

A novel model of complete microglial deficiency in the mouse arose from germ-line deletion of a super enhancer in the mouse *Csf1r* locus, the Fms intronic regulatory element (FIRE) *Csf1r*^ΔFIRE/ΔFIRE^ (*Fireko*) mice lack microglia throughout development^23^. Based upon analysis of these mice, many proposed functions of microglia in brain development appear redundant^24,25^ (and reviewed in ^26^). Instead, the congenital absence of microglia is associated with increased sensitivity/acceleration in models of neuropathology^27,28^.

The rat has many advantages and differences compared to the mouse as a model for the study of brain development and behaviour^29,30^. There are also major differences in biology of the mononuclear phagocyte system^7,31^. There have been relatively few studies of microglial function in rat development. We previously generated a *Csf1rko* rat and analysed the impact on the transcriptomic profile of multiple brain regions^12,32–34^. However behavioural and cognitive analysis of these animals was compromised by ventricular enlargement in the brain and growth retardation, developmental delay and early mortality. Here we describe the generation and characterisation of a *Csf1r*^ΔFIRE/ΔFIRE^ (*Fireko*) rat model of microglial deficiency.

## Results

### Generation of Csf1r^ΔFIRE/ΔFIRE^ (Fireko) rats

The rat FIRE sequence was identified by aligning mouse FIRE against the rat genome using the BLAST-like alignment tool (BLAT) on the UCSC genome browser (genome.ucsc.edu). Guide RNAs (gRNAs) designed to delete FIRE (Table 1) were validated in a rat kidney epithelial cell line (NRK-52E^35^). Cells were transfected with ribonucleoprotein (RNP) complexes containing pairs of gRNAs. One pair (109F/685R) resulted in the expected deletion (Supplementary Figure 1a) following sequence confirmation (data not shown). To produce *Fireko* rats, the RNP complex was electroporated into Long-Evans (LE) rat oocytes. One founder with the expected 585 bp deletion was bred to *Csf1r*^+/+^ LE rats. Heterozygous offspring from different mating pairs were bred together to produce *Fireko* rats. The frequencies of *Csf1r*^+/+^, *Csf1r*^+/ΔFIRE^ and *Fireko* rats at weaning were 26%, 51% and 23%, respectively (*n* = 389), not significantly different from Mendelian ratio (by contrast to 20-30% pre-weaning loss of *Fireko* mice^23^. Like the *Fireko* mice the homozygous mutant rats were indistinguishable from littermates at birth. There were no significant differences in body or major organ weights at 12 weeks of age between genotypes (Supplementary Figures 1b-c). Adult *Fireko* rats displayed increased porphyrin production around the eyes as seen in the *Csf1rko* rats^12^ but were not osteopetrotic and the bone marrow cavity, growth plate and trabeculae appeared normal (data not shown).

**Table 1.**
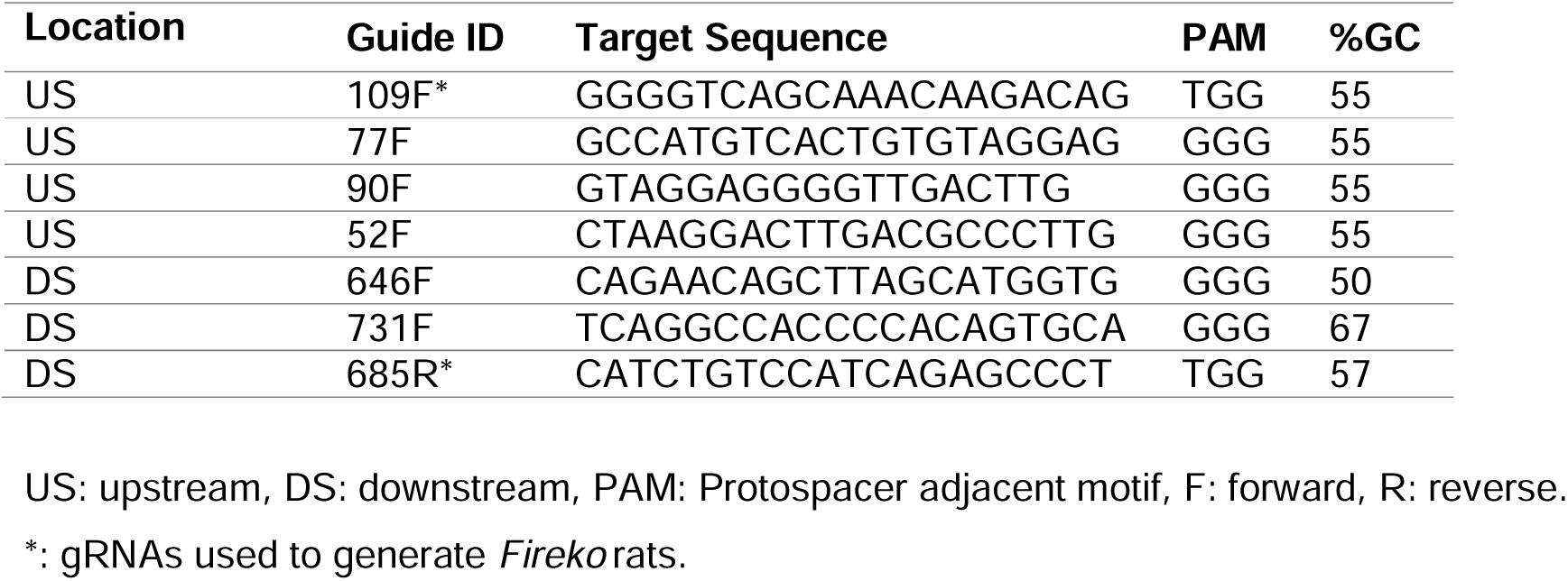
Candidate gRNAs.

### Monocyte populations in Fireko rats

*Fireko* mice retained monocytes in the blood and bone marrow (BM) but both classical LY6C^high^ and non-classical LY6C^low^ monocytes had almost undetectable CSF1R expression and did not respond to CSF1 *in vitro*^23^. This finding has been revisited recently; demonstrating that surface CSF1R in the non-classical monocytes is down-regulated by CSF1. Treatment of *Fireko* mice with blocking anti-CSF1R antibody leads selective depletion of Ly6C^low^ monocytes^36^ as seen previously in wild type mice^37^. Monocyte subsets in rats can be identified using antibodies against sialophorin (CD43) and HIS48^38^. We found that FIRE deficient rats had similar percentages of circulating granulocytes and total monocytes but the proportion of monocytes subsets differed to littermate controls. There was an increase in CD43^+^ non-classical monocytes and a decrease in CD43^−^ monocytes (Figure 1a). The full gating strategy is shown in Supplementary Figure 2. Fluorescently labelled porcine CSF1-Fc^39^ was used as a surrogate marker for CSF1R expression as our previous studies in *Csf1rko* rats showed complete loss of binding in the absence of *Csf1r*^12^. Whilst CSF1R expression detected by ligand binding was unaffected in CD43^+^ monocytes, it was reduced in a dose-dependent manner in the CD43^−^ monocytes of heterozygous and homozygous rats (Figure 1b).

**Figure 1-.**
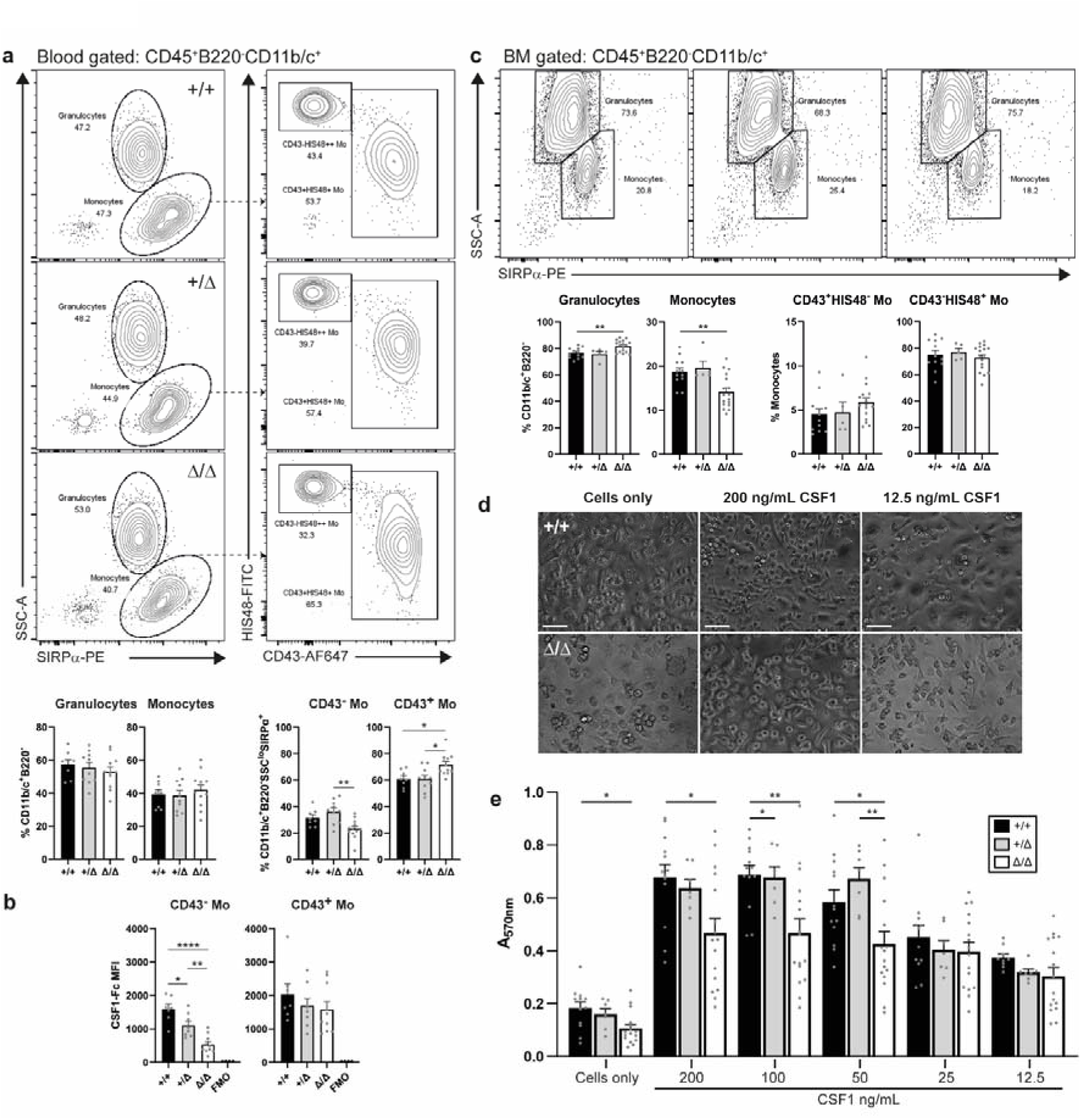
The affect of FIRE deletion on rat blood and bone marrow monocyte populations. **a)** Whole EDTA-blood from 2–3-month-old rats was analysed by flow cytometry. Live single cells were gated CD45^+^SIRPα^+^B220^−^ (Supplementary Figure 2) to identify granulocytes (SIRPα^+^SSC^high^) and total monocytes (SIRPα^+^SSC^low^) which were then divided into CD43^+/−^monocyte (Mo) populations. *n* = 8+/+, 10+/Δ and 12 Δ/Δ. *P* = 0.0014 (**), 0.0166 (*+/+ vs. Δ/Δ) and 0.0101 (*+/Δ vs. Δ/Δ). **b)** The mean fluorescence intensity (MFI) of porcine CSF1-Fc binding was determined on CD43^+/−^ monocyte subsets identified above. *n* = 7+/+, 8+/Δ and 10 Δ/Δ. *P* = 0.0233 (*), 0.0044 (**) and <0.0001 (****). **c)** Bone marrow (BM) from 2–4-month-old rats was analysed by flow cytometry. The gating strategy was the same as for the blood above. *n* = 12+/+, 5+/Δ and 17 Δ/Δ. *P* = 0.0076 (** granulocytes) and 0.0050 (** monocytes). **d)** Representative images of BM-derived macrophages after 7 days in culture. Scale bar = 50 µm. **e)** BM cells were put in culture in media alone or various concentrations of recombinant human CSF1. A MTT assay was performed on day 7. *n* = 7 – 16 rats per genotype. *P* = 0.0107 - 0.0426 (*) and 0.0035 - 0.0079 (**). All graphs show mean + SEM and *P* values were determined by one-way ANOVA with Tukey’s multiple comparisons test. Each data point represents an individual animal.

We also examined the monocyte populations in the BM by flow cytometry. The full gating strategy is shown in Supplementary Figure 2. There was a slight decrease in the percentages of CD11b/c^+^SIRPα^+^ monocytes that was coupled with an increase in granulocytes. Unlike the blood, there was no difference in the percentages of monocyte subsets based on CD43/HIS48 expression (Figure 1c). We cultured BM cells from rats either alone, or in a series of CSF1 concentrations for 7 days before using MTT to assess cell metabolic activity. Cells cultured in media alone differentiated into macrophages as unlike mice, rat monocytes express CSF1^40^ (Figure 1d). Differentiation was increased in the presence of added CSF1. When comparing genotypes, there were significant decreases in *Fireko* BM, but only at the higher concentration of CSF1 (Figure 1e). We also examined the BM-derived macrophages (BMDM) via flow cytometry. BMDM from all 3 genotypes expressed the same cell surface markers at the same levels. CSF1R expression (as determined by binding of labelled CSF1-Fc) was also reduced in the *Fireko* BMDM (Supplementary Figure 2).

### Macrophage populations in Fireko rats

*Csf1rko* rodents have a widespread loss of tissue macrophages^8,12^, whereas *Fireko* mice only lacked brain microglia and resident macrophages in select tissues^23^. We used a combination of immunohistochemistry and flow cytometry to examine the same tissue macrophage populations in *Fireko* rats. We began by examining 3 populations that were lost in the FIRE deficient mice: kidney, cardiac and large peritoneal macrophages. The full gating strategies are shown in Supplementary Figure 3.

In mice, the two largest cardiac macrophage populations are CD11b^+^ and are high or low in MHCII expression^41^. These populations were both reduced in FIRE deficient mice^23,42^. *Fireko* rats did not have a reduction in total cardiac myeloid cells by flow cytometry, but rather the types of SIRPα^+^CD11b/c^+^ cells differed. The knockout rats had a reduction in the MHCII^++^ resident macrophage population whilst the proportion of CD43^+^ monocytes increased. There was no change in the percentages of total granulocytes, CD43^−^ monocytes or MHCII^+^ macrophages (Figure 2a). Kidneys from the same cohort were analysed. The resident kidney macrophage population in mice is CD11b^mid^ ^43^ and this population was lost in rats. As in the heart, there was an increase in the CD43^+^ monocytes (Figure 2b).

**Figure 2-.**
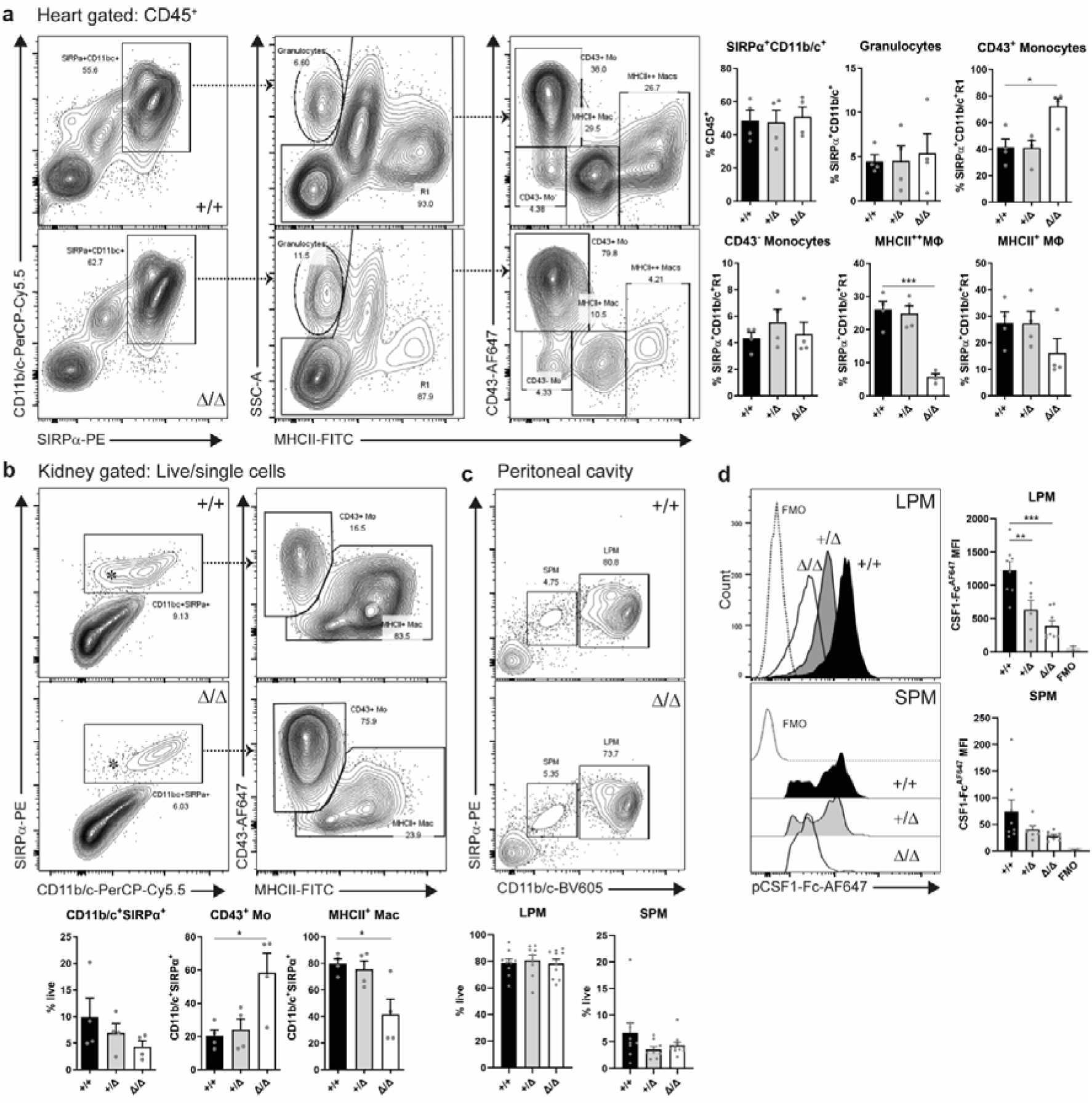
FIRE deletion in rats results in a loss of cardiac and kidney macrophages, but not large peritoneal macrophages. Male and female rats aged 7-11 weeks were perfused with PBS and fixed single cell suspensions of enzymatically digested hearts (a) and kidneys (b) were analysed by flow cytometry. **a)** Representative flow cytometry profiles of hearts gated on CD45^+^ cells. *n* = 4 rats per genotype. *P* = 0.0126 (*) and 0.0002 (***). **b)** Representative flow cytometry profiles of kidneys gated on live/single cells. *n* = 4 rats per genotype. *P* = 0.0219 (* Mo) and 0.0217 (* Mac). *: CD11b/c mid cells, Mo: monocyte, Mac/MΦ: macrophage. **c)** Peritoneal lavages from male and female rats aged 10-12 weeks were analysed by flow cytometry. Representative flow cytometry profiles show small peritoneal macrophages (SPM, CD11b/c^+^SIRPα^+^) and large PM (LPM, CD11b/c^++^SIRPα^++^). Cells were gated on live, single cells. *n* = 9-10 rats per genotype. **d)** The mean fluorescence intensity (MFI) of pCSF1-Fc^AF647^ binding in SPM and LPM. *n* = 6-8 rats per genotype. *P* = 0.0069 (**) and 0.0001 (***). FMO: fluorescence minus one. Graphs show mean + SEM and *P* values were determined by one-way ANOVA with Tukey’s multiple comparisons test. Each data point represents an individual animal.

There are two populations of peritoneal macrophages (PM) that can be distinguished by their size. Large PM (LPM) are the most abundant and are CD11b^++^F4/80^++^ in mice whereas small PM (SPM) express lower levels of these markers^44^. In rats, these same populations can be identified using antibodies against SIRPα (CD172a) and CD11b/c^12^. Whilst the large PMs were absent in *Fireko* mice^23^, these were retained in the rats (Figure 2c). The remaining PM in *Fireko* mice had completely lost CSF1R expression with a partial loss observed in heterozygotes^23^. Unlike the mice, *Fireko* rats did not have a complete loss of CSF1R in either the small or large PMs (Figure 2d) but levels were reduced compared to littermate controls. Whilst the LPM showed uniform binding of labelled CSF1, the SPM population was more heterogenous. This is likely due to the various levels of CSF1R observed in rat PMs using the *Csf1r*-mApple transgene^38^.

Macrophage populations in the liver and spleen were unaffected by FIRE deletion in the rat (Supplementary Figure 4). In overview, deletion of FIRE has a less penetrant impact on CSF1R expression and tissue macrophages in rats than previously reported in mice.

### Fireko rats are microglia deficient

*Csf1rko* rats on an inbred background develop substantial ventricular enlargement, involution of the olfactory bulb and disruption of the corpus callosum by 3 weeks of age^33^. By contrast, in adult *Fireko* rats the brain-body weight ratio was unchanged compared to littermate controls (Supplementary Figure 1b) and there were no evident structural abnormalities in the majority of animals (Figure 3a). We did observe ventricular enlargement in some rats, likely due to genetic differences of the outbred background strain (Long Evans). Importantly, none of the rats had to be culled for welfare concerns, even when aged to 11 months. Staining for IBA1 demonstrated the complete absence of microglia in the forebrain (Figure 3b). There were few IBA1^+^ perivascular macrophages within the parenchyma surrounding the hippocampus. The reduction of these CD11b/c^low^CD45^high^ cells and loss of microglia (CD11b/c^high^CD45^low^) was confirmed by flow cytometry. There was no alteration in microglial density in the heterozygotes (Figure 3c). The full gating strategy is shown in Supplementary Figure 5a. The same selective loss of microglia was observed in the retina (Supplementary Figure 5b). Within the choroid plexus, stromal macrophages were present, but epiplexus macrophages were not detected (Supplementary Figure 5c), as in the *Fireko* mice^45^. Meningeal macrophages were present in wild type rats but were obviously reduced in *Fireko* rats (Supplementary Figure 5d). Aged *Fireko* mice develop calcium deposits and axonal spheroids, hallmarks of CSF1R-related leukoencephalopathy in patients^46^. Both forms of pathology were also detected in *Fireko* rats aged 9-10 months and were predominantly located in the thalamus (Figure 3d-e).

**Figure 3.**
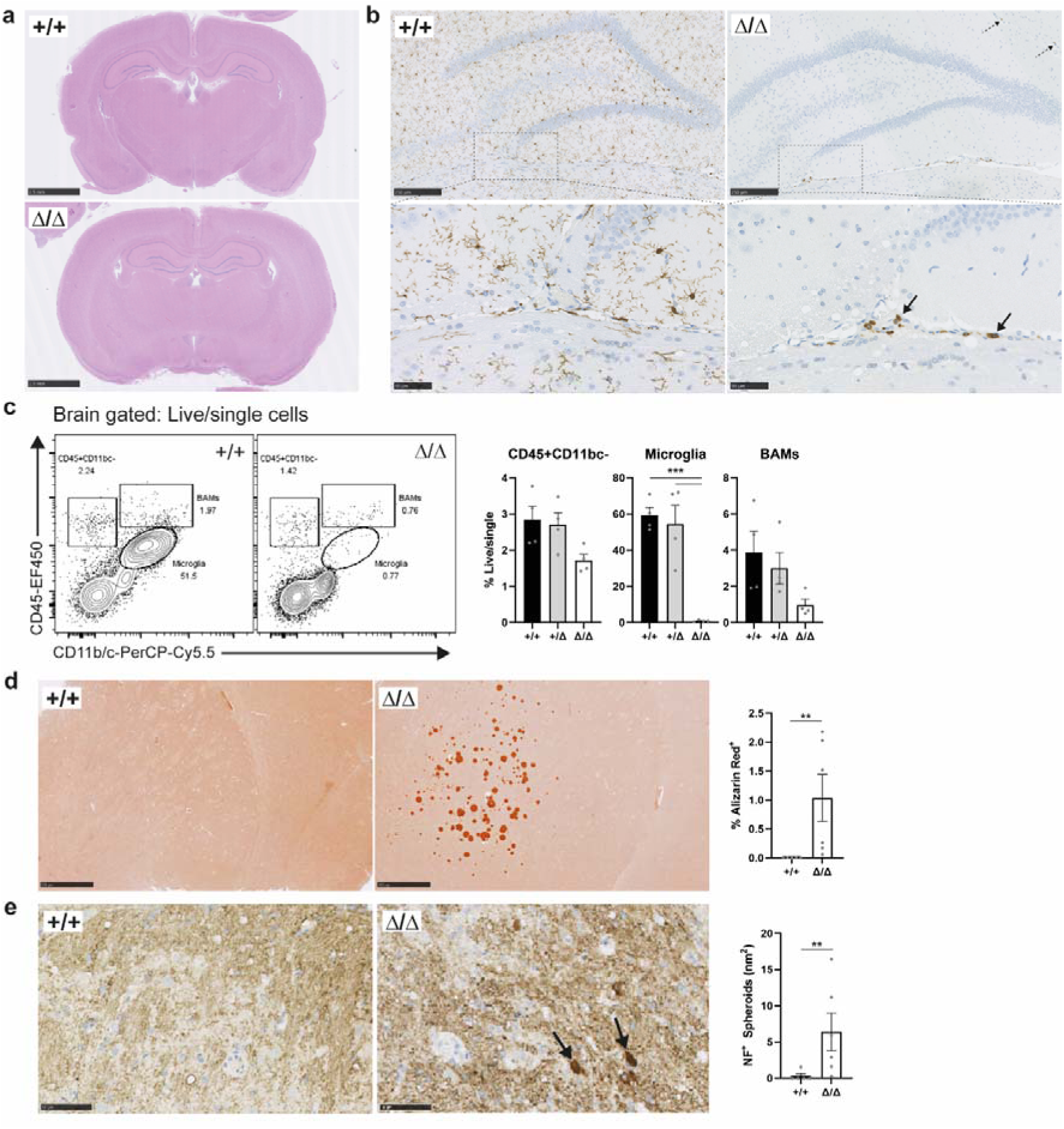
FIRE deletion in rats results in a loss of microglia leading to calcification and axonal spheroids. **a)** H&E staining of adult rat brains. Scale bars: 2.5 mm. Images are representative of 4 rats per genotype. **b)** Paraffin-embedded formalin-fixed adult brains aged 2-3 months were stained with an antibody against IBA1. Dotted arrows: perivascular macrophages in the hippocampus. Solid arrows: monocytes. Images representative of 5 +/+ and 6 Δ/Δ. Scale bars: 250 µm and 50 µm (inset). **c)** Single cell suspensions from myelin-depleted of brains from adult rats aged 2-3 months were analysed by flow cytometry. Microglia: CD45^low^CD11b/c^+^. Border associated macrophages (BAM): CD45^+^CD11b/c^+^. Graphs show mean + SEM and *P* values were determined by one-way ANOVA with Tukey’s multiple comparisons test. *** *P* = 0.0004 (+/+ vs. Δ/Δ) and 0.0007 (+/Δ vs. Δ/Δ). *n* = 4 rats per genotype. **d)** and **e)** Paraffin-embedded formalin-fixed adult brains aged 9-10 months were stained with Alizarin Red S to detect calcium (d) and an antibody against neurofilament (NF) to detect axonal spheroids (e). Arrows point to NF^+^ spheroids. Scale bars: 500 µm (d) and 50 µm (e). Images representative of 5 +/+ and 6 Δ/Δ. Graphs show mean ± SEM and *P* values were determined by Mann-Whitney U test. ** *P* = 0.0043 (calcium) and 0.0087 (spheroids). Each data point represents an individual animal.

To assess the impact of microglia loss on other cell types, we performed RNA-seq on the hippocampus and striatum of adult *Fireko* and littermate controls aged 11 weeks. The data are provided in Supplementary Table 1. The expression data were subjected to network analysis using *Graphia* as previously used for the *Csf1rko* rats^12^. The clusters are presented in Supplementary Table 2. The largest set of transcripts (Cluster 8, Figure 4a) that showed a clear relationship to genotype includes *Csf1r* and 221 others including *Aif1*, *C1qa* and *P2ry12* (Figure 4b). This set overlaps but is significantly larger than the 105-transcript microglial signature defined by analysis of 3-week-old *Csf1rko* rats^32^. There were two smaller clusters related to genotype containing macrophage-associated genes. Cluster 27 (30 transcripts) includes *Cd68, Cd74* and Class II MHC genes. Cluster 164 (8 transcripts) includes *Cd163,* and *Mrc1* (CD206). These clusters were separated from the main microglia-associated cluster because the transcripts were not down-regulated to the same extent and/or their abundance differs between hippocampus and striatum. Aside from *Csf1r*, reduced by around 50% consistent with the lack of dosage compensation^12^, none of the transcripts in these clusters was altered to the same extent in heterozygous rats (Figure 4b). Notably, we detected no significant difference in abundance of neuronal transcripts that distinguish hippocampus and striatum, nor markers of neurogenic progenitors (*Dcx*), oligodendrocytes (e.g. *Olig1, Mbp*) or astrocytes (*Gfap*) indicating that apart from the loss of microglia, the overall cellular composition and differentiation is unchanged. In *Csf1rko* rats^32^, the stress response gene *Rbm3* was increased in all brain regions but this was not significantly increased in *Fireko* brains.

**Figure 4-.**
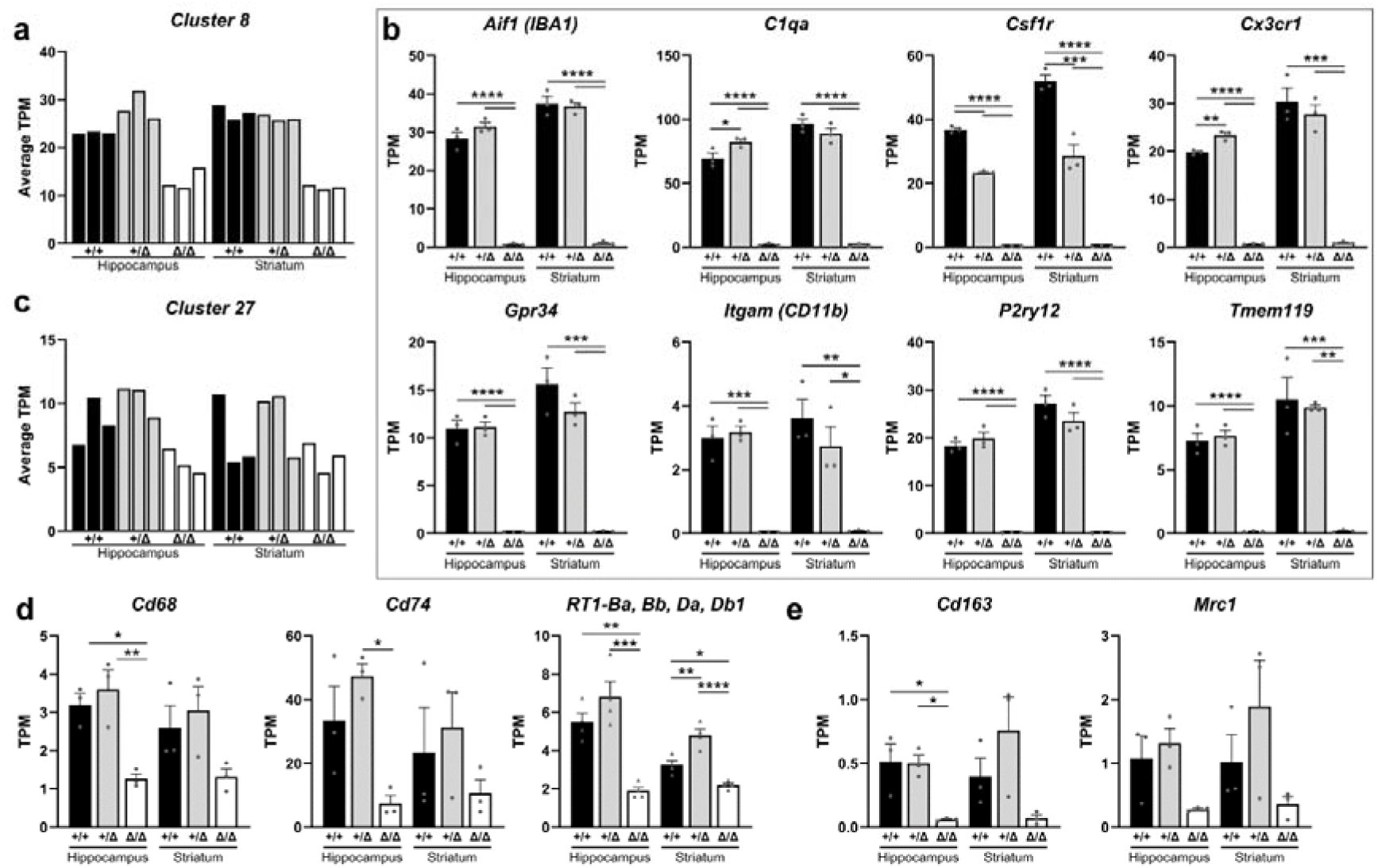
The effect of FIRE deletion on gene expression in the hippocampus and striatum. RNA-seq analysis was performed on hippocampus and striatum from *Fireko* and littermate controls aged 11 weeks (*n*=3/group). Gene co-expression analysis was performed using Graphia as described in the methods. The full data set and clusters lists are provided in Supplementary Tables 1 and 2. **a)** Average expression profile of co-expressed genes in Cluster 8 (222 genes). **b)** Gene expression profiles for individual selected microglia-associated genes from Cluster 8. Graphs show mean + SEM and *P* values were determined by one-way ANOVA with Tukey’s multiple comparisons test. *P* = (*) 0.0371 *C1qa* and 0.0188 *Itgam*, (**) 0.0012-0.0049, (***) 0.0001-0.0009, (****) <0.0001. **c)** Average expression profile of co-expressed genes in Cluster 27 (30 genes). **d)** Gene expression profiles for individual selected macrophage-associated genes from Cluster 27. RT1 = MHC Class II genes. Graphs show mean + SEM and *P* values were determined by one-way ANOVA with Tukey’s multiple comparisons test. *P* = (*) 0.0135-0.0314, (**) 0.0030-0.0073, (***) 0.0003, (****) <0.0001. **e)** Gene expression profiles for individual selected macrophage-associated genes from Cluster 164. Graphs show mean + SEM and *P* values were determined by one-way ANOVA with Tukey’s multiple comparisons test. *P* = (*) 0.0244 (+/+ vs. Δ/Δ) and 0.0282 (+/Δ vs. Δ/Δ). Each data point represents an individual animal.

**Table 2.**
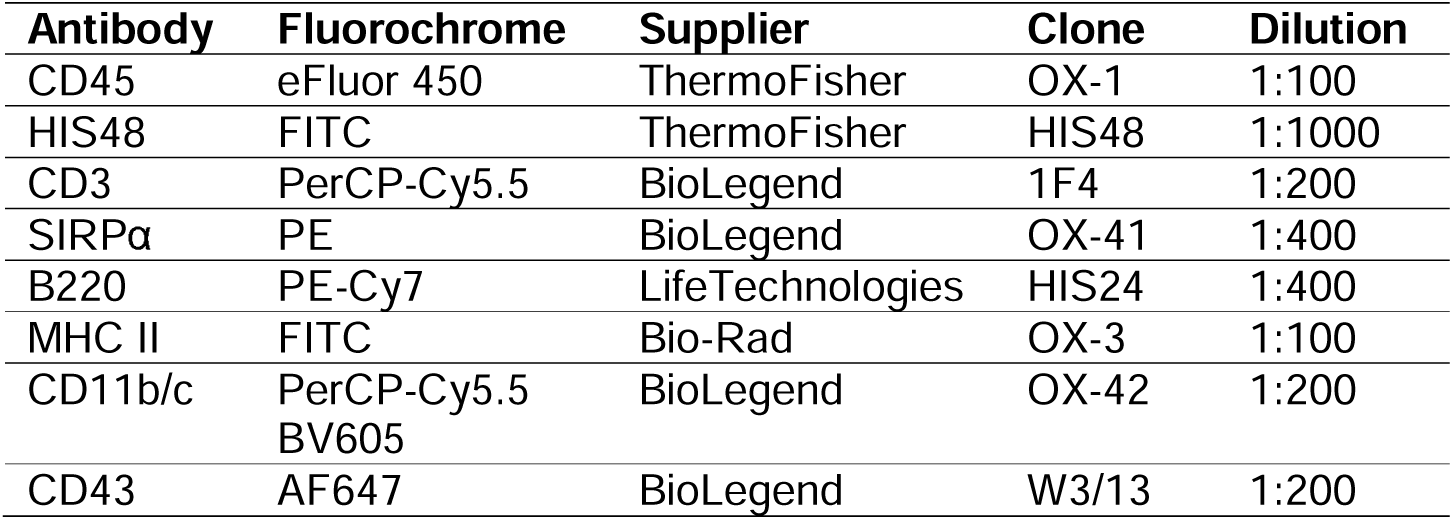
Antibodies used for flow cytometry.

### Ethologically relevant behaviours are unaffected by loss of microglia

Microglia have been proposed as key regulators of adult neurogenesis^47,48^ although the gene expression analysis provided no evidence of a deficiency in pathway in the *Fireko*. Previous research has indicated that hippocampal adult neurogenesis is greatly enriched in rats compared to mice and contributes to performance in behavioural tasks^30^. We performed a suite of behavioural tests on *Fireko* rats and their littermate controls to investigate the potential impact of microglia loss on behaviour (Figure 5a and Supplementary Methods).

**Figure 5-.**
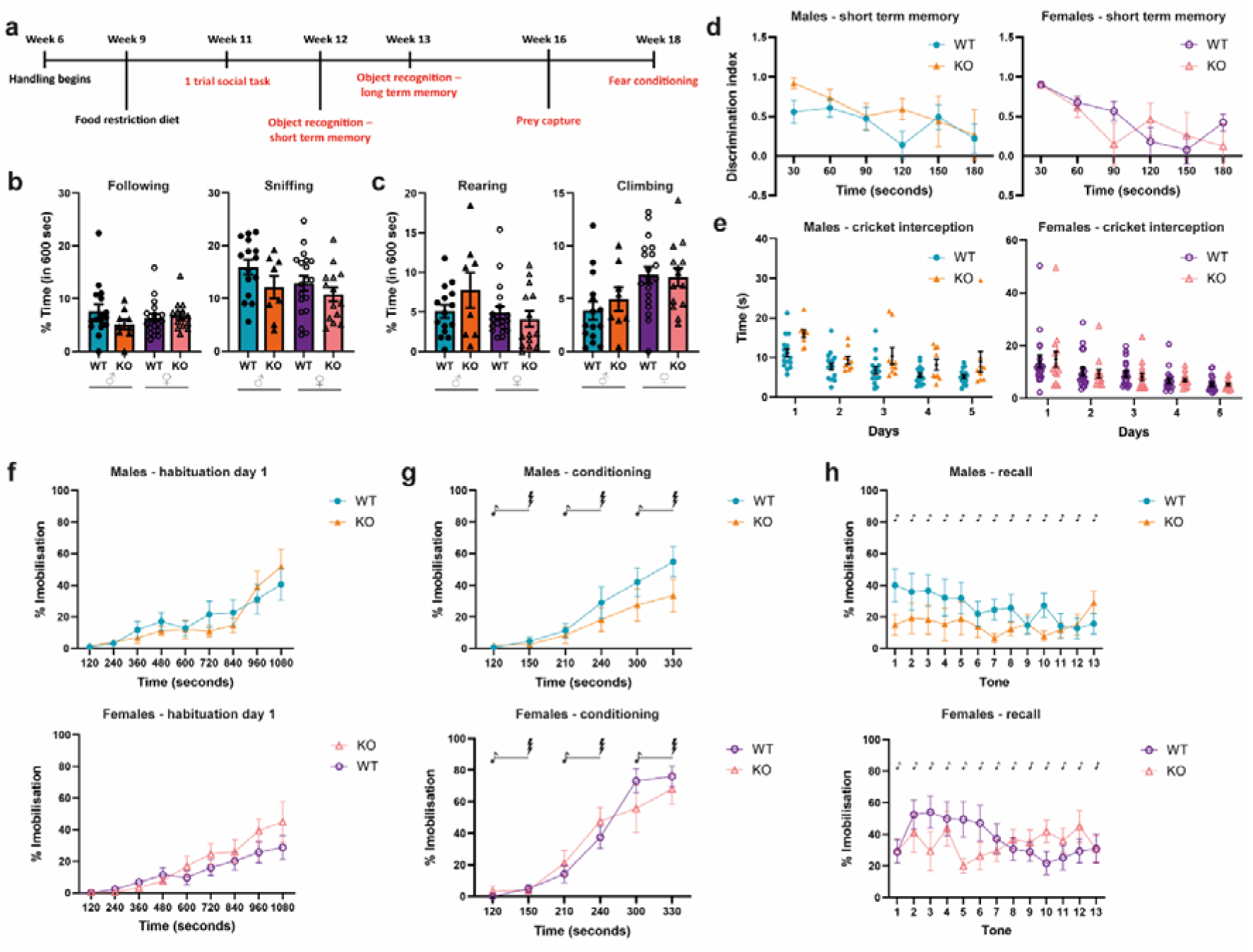
Loss of microglia had no effect on various behaviours in the rat. **a)** Timeline of behavioural testing pipeline for FIRE rats. **b)** Social behaviours of 11 week old rats during a one trial social interaction task. **c)** Exploratory behaviours recorded during the same task. **d)** Short-term object recognition memory in 12 week old male and female rats. **e)** Prey capture performance of 16 week old male and female rats. **f** - **h)** Fear conditioning in 18 week old male and female consisting of habituation (f), conditioning with tone-shock pairings (g) and fear recall (h). All graphs show mean ± SEM and *P* values were determined by two-way ANOVA with Tukey’s multiple comparison test. All *P* values > 0.05. *n* = 9-19 rats per genotype and sex. Sexes were separated for illustrative purposes. WT: *Csf1r*^+/+^/ *Csf1r*^+/ΔFIRE^, KO: *Csf1r*^ΔFIRE/ΔFIRE^. Symbols: foot shock (lightning bolt) and tone (music note).

Rats are social, highly exploratory animals that live in social hierarchies and can distinguish between familiar and unfamiliar conspecifics (reviewed in ^49^). To determine the effect of microglia loss on social behaviours, we used a 1 trial social task involving exposure to an unfamiliar age- and sex-matched stimulus rat. Across all groups, regardless of sex or genotype, rats spent an average of 5-7% of the trial following the stimulus rat and showed comparable total time spent sniffing the unfamiliar conspecific, indicating similar levels of social interest (Figure 5b). Patterns of investigation, including a predominant focus on the genital area, also did not differ between groups when sniffing behaviour was broken down by body region (Supplementary Figure 6a). Non-social exploratory behaviours were also assessed by analysing exploration of the arena. Measures of anxiety-related behaviours including time spent wall climbing and rearing were comparable across all groups, with no distinction between *Fireko* rats and their littermate controls (Figure 5c).

To evaluate potential cognitive effects of microglia loss on hippocampus-dependant recognition, memory was assessed using standard object recognition tests. Short term memory was tested by presenting a novel object 5 minutes after exposure to two identical objects, while long-term memory was evaluated following a 24-hour delay. In both paradigms, *Fireko* rats demonstrated intact memory performance, with no deficits observed relative to littermate controls (Figure 5d and Supplementary Figure 6b).

Prey capture tasks provide a sensitive assay of sensorimotor integration and plasticity because successful performance requires precise coordination between sensory input (cricket) and motor output (capture). With repeated exposure, predation efficiency improves and eventually becomes a learned response (reviewed in ^50^). All rats showed improved performance over time, as evidenced by a decrease in the time taken to intercept the cricket over the course of the experiment, and this learning was unaffected by loss of microglia (Figure 5e). Similarly, no significant differences were observed between the *Fireko* rats and littermate controls in the time taken to catch and consume the cricket (Supplementary Figure 6c).

To assess associative learning and memory, auditory fear conditioning was used as the final behavioural paradigm. During habituation, all groups displayed similar baseline levels of immobility in the testing context, with immobility increasing over time (Figure 5f and Supplementary Figure 6d). During conditioning, immobility progressively increased over the course of 3 tone-shock pairings in all animals (Figure 5g), indicating comparable acquisition of the conditioned fear response regardless of sex or genotype. 24 hours after conditioning, all rats froze in response to the first presentation of the unreinforced conditioned tone; this conditioned fear response gradually diminished over repeated tone presentations (Figure 5h). A smaller cohort of rats (*n* = 4-9 per genotype and sex) also underwent auditory fear conditioning at 10 months of age and no significant differences were seen (data not shown).

## Discussion

Microglia-deficient *Fireko* mice have been studied in detail in order to understand the functions of microglia in brain development and age-dependent neuropathology and as a model for CSF1R-related leukoencephalopathy^23,25,51^. Here we have generated and analysed *Fireko* rats. The effects of the enhancer deletion in rats are subtly different from those reported in mice. The differences may not be entirely due to the difference in species. We purposely created the rat mutation on an outbred background to avoid possible complications seen in inbred mice. The enhancer deletion in *Fireko* rats appears to have a less penetrant effect on CSF1R expression than the same mutation in mice. Bone marrow cells remained responsive to CSF1, the receptor was more highly expressed in blood monocytes and peritoneal macrophages were retained where they are absent in *Fireko* mice^23^.

In both species, the homozygous FIRE deletion has a selective effect on the expression of the receptor on the surface of non-classical monocytes and shifts the ratio of subsets in favour of the non-classical. In rats, CD43^−^ non-classical monocytes are the dominant monocyte population^38^ and are selectively lost in *Csf1rko* animals^33^. Analysis of a knock-in CSF1R reporter in mice indicates that non-classical monocytes express higher levels of the receptor^52^. There is evidence in mice that the classical monocytes act as a sink for CSF1 and thereby regulate the differentiation and survival of non-classical monocytes which are absolutely CSF1R-dependent^53^.

The differences between mouse and rat may be related to discordant regulation of the ligand, CSF1, which is expressed by monocytes and macrophages in rats and other large animals (including humans) but not in mice^40^. The differentiation of macrophages from bone marrow in the absence of added growth factor (Figure 1d-e) suggests that rats may also have differential expression of other factors such as FLT3 ligand (*Flt3lg*) or GM-CSF. FLT3 was recently shown to be essential for monocytosis in humans but not in mice^54^. *Csf1r* was expressed normally in *Fireko* mouse tissues where macrophage numbers were not affected^23^. These include liver, lung, intestine and spleen and was attributed to the presence of multiple shadow enhancers in the *Csf1r* locus. FIRE is almost perfectly conserved between mouse and rat and the mouse FIRE sequence is capable of driving identical transgene expression in both species^38,55^. However, there is considerable sequence divergence in other areas of the *Csf1r* locus. It is therefore likely that there are species-specific shadow enhancers that partly explain the difference between the two rodents.

Like the mouse counterpart, *Fireko* rats appear entirely microglia deficient. Recent evidence in the mouse indicates that microglia can be replaced by monocyte-derived cells following depletion^56^. Despite the apparent retention of receptor activity in *Fireko* rat blood monocytes, microglia remained deficient after 11 months based on IBA1 staining. The complete loss of the microglia transcriptional signature was shared with previous analysis of microglia-deficient *Csf1rko* rats^32,34^. However, many genes that are strongly enriched in isolated microglia compared to total brain are not significantly depleted in the absence of microglia in *Fireko* mice^25^ or *Csf1rko* rats^32^. They include the key growth factor, *Igf1*, the TGFb regulator *Nrros1*, the commonly-used neuroinflammation marker *Tspo* and the complete set of lysosome-associated genes (reviewed in ^26^). The lysosomal gene, *Hexb*, has been considered a microglia-specific marker in the mouse^57^, but *Hexb* is reduced by less than 50% in *Fireko* rats. These observations suggest that other cell types can induce expression of key regulators to compensate for the absence of microglia.

*Fireko* mice develop an age-dependent neuropathology that has been considered a model for human CSF1R-related leukoencephalopathy^46,51^. Given their greater longevity, such an analysis in rats would probably require a cohort aged for 2 to 3 years. However, we did observe calcification in the thalamus and evidence of axonal spheroids at 9-10 months (Figure 3d-e). Neuroinflammatory markers such as *C4b* and *Serpina3n* were already increased in multiple brain regions in six weeks old *Fireko* mice^25^ but this was not evident in the RNA-seq analysis of the *Fireko* rat at 11 weeks.

Microglia are widely considered to be essential for normal postnatal brain development. Experimental deletion of microglia in mice was reported to lead to robust depletion of oligodendrocyte precursors and myelin-associated proteins^58^ but analysis of *Fireko* mice indicated that microglia are not essential for development myelination^24^. Similarly, we observed no structural impact on the corpus callosum in *Fireko* rats and expression of transcripts associated with myelination was unaffected. The proposed role of microglia experience-dependent plasticity of neuronal function was also not supported by analysis of microglia-deficient *Fireko* mice^25^. Here we extended those observations to a detailed analysis of multiple behavioural traits in *Fireko* rats. We found no significant difference between *Fireko* rats and littermate controls in any of the assays. It is of course possible that subtle differences would emerge with much larger cohorts and or with inbred lines to reduce the inter-individual variation. Any such analysis would need to consider the possible linked variation on the same chromosome as *Csf1r.* In the interim we can reasonably conclude that microglia are not absolutely required for normal postnatal neurodevelopment in either rats or mice.

Microglia clearly do participate in developmental and homeostatic clearance functions in the brain. In interpreting the effect of the *Fireko* it is important to recognise that congenital microglia deficiency is quite distinct from experimental microglial depletion or mutations in microglia-expressed genes. We suggest that dying or dysfunctional microglia are likely to interfere with the ability of other cells to compensate for the loss of microglial functions^26^. The identity of the cells and mechanisms that enable relatively normal development without microglia remain to be determined.

## Methods

### Cell culture

NRK-52E cells (ATCC^®^ CRL-1571™)^35^ were obtained from Laura Denby (University of Edinburgh) and cultured in DMEM supplemented with 10% fetal calf serum (FCS) and 2 mM GlutaMAX (Life Technologies). Cells were passaged at a 1:10 dilution every 48-72 h (at confluence) with TrypLE™ Express (Gibco) and maintained at 37°C, 5% CO_2_.

### CRISPR design and validation

gRNAs were designed to complement the 200 bp flanking sequences of rat FIRE (chr18:56,434,670-56,435,007). The CRISPOR website^59^ was used with the following selection criteria: “Rattus norvegicus – Rat – UCSC Jul. 2014 (RGSC 6.0/rn6)”, “PAM: 20 bp-NGG – Sp Cas9, SpCas9-HF1, eSpCas9 1.1”. gRNAs were chosen to have: GC content 50-70%, a “G” immediately upstream the PAM site at the 3’-end, a “G” at the 5’-end and off target effects minimized^60^. The resulting gRNAs are outlined in Table 1.

The assembly of ribonucleoprotein (RNP) complexes and subsequent transfection into NRK-52E cells was performed using reagents from Integrated DNA Technologies (IDT) unless specified otherwise. Methods were according to instructions (Alt-R CRISPR-Cas9 system, IDT). In brief, single gRNAs were assembled into RNP complexes and pairs containing an upstream and downstream gRNA were then transfected into NRK-52E cells (1.5×10^5^ cells/mL) using Lipofectamine CRISPRMAX reagent and Opti-MEM media (Thermo Fisher Scientific). DNA was purified from transfected cells by digesting in 100 mM Tris HCl pH8.5, 0.2% SDS, 200 mM NaCl, 5 mM EDTA and 400 µg/mL Proteinase K (QIAGEN) at 55°C for 1 h followed by standard phenol/chloroform purification. Conventional PCR was performed using Q5 high-fidelity polymerase (New England Biolabs) and the primers F2-5’:

TCTCCTCTGTTCTACATCCAGC and R1-5’:GCCAAGCCCTGACTTTTCAT at an annealing temperature of 61.5°C. PCR products were run using the E-Gel™ Power Snap Electrophoresis Device using precast E-Gel™ CloneWell II gels (Invitrogen) to obtain 1,051 bp (*Csf1r*^+/+^) or 466 bp (*Fireko*). PCR products ∼ 500 bp were collected for a nested PCR using Q5 polymerase (as above) using the primers F1-5’: GCAGACATTAAACCAGGCTGA and R2-5’: TCCTGCTGTCCCATTCCTTT. PCR amplicons were separated by standard electrophoresis, purified with the MinElute Gel Extraction Kit (QIAGEN) and Sanger-sequenced using the same primers.

### Generation of Csf1r^ΔFIRE/ΔFIRE^ (Fireko) rats (LE-Csf1r^em1Prdns^)

The animal experiments underwent review by the University of Edinburgh Welfare and Ethical Review Body (AWERB). They were subsequently approved by the UK Animals in Science Regulation Unit (ASRU) under the Animals (Scientific Procedures) Act 1986 and authorised by Project Licence PP8165520 in strict accordance with the Home Office Code of Practice.

Rats were bred and housed under specific pathogen-free conditions. *Fireko* rats were produced by electroporation of 2-day old oocytes with the RNP complex containing the gRNAs 109F/685R. Both donor and recipient females were Long-Evans (Charles River Crl:LE). Males and females were used in all analyses.

### Animals

All animals housed for experimental purposes were kept according to the UK Animals (Scientific Procedures) Act 1986 following the Home Office Code of Practice. Rats were housed in Allen Town Next Gen 1800 IVC cages, at 12 air changes/h on Eco-Pure Aspen 2HK wood chip bedding, with Sizzle Nest nesting, and Maxi Plus Rat Tunnels (150×82×3 mm), Aspen Bricks (50×100×100 mm) (all from Datesand) as standardised environmental enrichment per cage. Water was supplied through an auto water system (Avidity). Rats were all fed RM3 (breeding) or RM1 (stock) pelleted diet (Special Diet Services). Rats were separated by sex at wean and housed in groups based on the cage to weight ratio of the caging. The environmental conditions were 20-24°C, 45-65% relative humidity with a 12 h on/off light cycle from 0700 to 1900 h. Animals were changed to fully cleaned bedding cages weekly which were autoclaved at 121°C (Vakulab PL201019-2G, MMM). Cages were washed prior at 80°C for 20 min wash/rinse cycles (Noedisher TP acid wash, Atlantis, Tecniplast). Animals were microbiologically screened quarterly according to FELASA recommendations. Our breeding program followed the guidelines outlined in the UFAW Handbook. Briefly, continuous breeding pairs were established at 8 weeks of age. Breeding pairs were typically replaced every six months or after the sixth litter is weaned.

### Preparation of single cell suspensions for flow cytometry

Blood was collected directly from the heart into EDTA tubes from rats under isoflurane and kept on ice. Blood was washed 3 times in PBS prior to staining with Zombie fixable viability dye (BioLegend) according to instructions.

Bone marrow (BM) isolation and peritoneal lavages were performed as previously described for mice^61^. BM was cryopreserved in FCS containing 10% DMSO and thawed 2 h prior to flow cytometry analysis. Cells were maintained at 37°C, 5% CO_2_ during this time in complete media: RPMI 1640 containing GlutaMAX™ (Invitrogen), Penicillin-Streptomycin solution (Gibco) and 10% FCS. Cells were cultured on non-tissue-culture treated petri dishes (Sterilin). The generation of BM-derived macrophages was achieved by culturing BM cells in 100 ng/mL recombinant human (rh) CSF1 for 7 days.

Organs were harvested from rats perfused with PBS (for heart, kidney and brain flow cytometry) or culled via a rising concentration of CO_2_ (spleen and liver flow cytometry) and finely minced with curved scissors in flow buffer (PBS +2% FCS) on ice. After centrifugation (400 *xg*, 5 min, 4°C), 1 mL of enzyme cocktail was added per 0.1 g of tissue. The cocktail was prepared in RPMI containing 0.625 mg/mL Collagenase D from *Clostridium histolyticum* (Merck), 0.425 mg/mL Collagenase V from *Clostridium histolyticum* (Merck), 1 mg/mL Dispase II (Life Technologies) and 150 µg/mL DNAse I grade II from bovine pancreas (Merck). Organs were incubated at 37°C in a shaking oven for 40 min (1 h for kidneys) then passed through 70 µm cell strainers moistened with flow buffer. Following centrifugation (400 *xg*, 5 min, 4°C), cell pellets from non-perfused animals were resuspended in 5 mL red blood cell (RBC) lysis buffer (BioLegend) and incubated on ice for 5 min in the dark. Flow buffer (15 mL) was added before another centrifugation step. Cell pellets were then resuspended in flow buffer for same-day analysis or washed 3 times in PBS prior to staining with Zombie fixable viability dye (BioLegend) according to instructions.

Single cell suspensions of myelin-depleted brains from PBS perfused rats were prepared according to^12^.

### Flow cytometry

Single cell suspensions were incubated with mouse anti-rat CD32 clone D34-485 (Fc block, BD Biosciences) at 1 µg/100 µL cells and incubated on ice for 20 min. Primary antibodies (Table 2) were added and cells incubated for 20 min on ice, in the dark. After a wash step with flow buffer, cells were either directly analysed on a BD Fortessa (spleen, liver, peritoneal lavages) with dead cells excluded with DAPI or fixed for analysis the following day (heart, kidney, brain).

For fixation, cells were resuspended in 50 µL PBS then 50 µL of 2% paraformaldehyde was added. After a 15 min incubation in the dark at RT, cells were centrifuged at 400 *xg* for 5 min, resuspended in 150 µL flow buffer, then centrifuged again. Samples were resuspended in 400 µL flow buffer and kept at 4°C overnight.

### Bone marrow macrophage viability assay

Cryopreserved BM cells from adult rats were plated at 2.5×10^5^ cells/well of a 96-well plate in complete media or various concentrations of rhCSF1. Plates were incubated at 37°C, 5% CO_2_ for 7 days. Supernatants were removed and 50 µL of 3-(4,5-dimethylthiazol-2-yl)-2,5-diphenyltetrazolium bromide (MTT from Merck; 1 mg/mL, diluted in sterile PBS) was added. After a 3 h incubation the MTT was removed and 100 µL was isopropanol added to solubilise the purple formazan for 10 min at 37°C, 5% CO_2_. The plates were read at 570 nm using a Synergy HT reader.

### Immunohistochemistry

Formalin-fixed paraffin embedded organs were stained as previously described^12^ with antibodies against IBA1 (rabbit polyclonal, Biosensis, 1:2000) and Neurofilament (clone SMI 312, BioLegend, 1:4000). Antigen retrieval was performed using citric acid-based antigen unmasking solution (Vector Laboratories) in a microwave, boiling for 5 min.

### Calcium staining

A 2% Alizarin Red S (Merck) solution was prepared in dH_2_O and the pH adjusted to 4.1 – 4.3 with 10% ammonium hydroxide (Thermo Scientific Chemicals). Formalin-fixed paraffin embedded organs were stained for 1 min after being dewaxed and rehydrated. Slides were dehydrated in acetone (20 dips), acetone:xylene (1:1, 20 dips) and cleared in xylene (2x 5 min) before mounting.

### Image acquisition and quantification

Whole-slide brightfield images were acquired using the NanoZoomer slide scanner. Image analysis was performed with NDP.view2 software v2.9.29 (Hamamatsu) and ImageJ v1.54p. For liver and spleen, 10 regions of interest at 20X magnification were exported per sample as .tiff files from the NDP.view files. IBA1^+^ signal was quantified from the whole area corresponding to each ROI (i.e., each tiff file). For calcium staining (2.5X) and spheroid detection (10X), 1 and 3 .tiffs were exported from the thalamus, respectively.

### Bulk RNA-seq

The striatum and hippocampus from male and female rats aged 11 weeks were snap frozen in dry ice. RNA was isolated as in ^12^. RNA-seq libraries were prepared by Edinburgh Genomics, The University of Edinburgh, with the Illumina Stranded mRNA-seq ligation protocol. Sequencing of 18 samples was performed using a single NovaSeq S1 100 cycle sequencing run on a NovaSeq 6000 machine (Illumina). Sequencing depth was between 98 million and 133 million paired end reads per sample. The raw sequencing data, in the form of fastq files, are deposited in the European Nucleotide Archive under study accession number PRJEB94302.

### RNA-seq processing and analysis

Raw reads were pre-processed using fastp v0.23.2^62^ using previously described parameters^33^. FastQC^63^ was used on pre and post-trimmed reads to ensure adequate sequence quality, GC content, and removal of adapter sequences. The reference transcriptome used in this study was created by combining the unique protein-coding transcripts from the Ensembl and NCBI RefSeq databases of the Rnor6.0 annotation, as previously described^64^. After pre-processing, transcript expression level was quantified as transcripts per million using Kallisto (v0.46.0)^65^. Kallisto expression files were imported into RStudio (R version 4.2.1) using the tximport package (v1.24.0) to collate transcript-level TPM generated by Kallisto into gene-level TPMs for use by downstream tools.

### Network cluster analysis of gene expression

All network cluster analysis was conducted using Graphia (https://graphia.app)^66^. For RNA-seq data, only genes expressed at ≥ 1 TPM in at least 2 samples were retained for analysis. Graphia calculates a Pearson correlation matrix comparing similarities in expression profiles between samples, and only relationships where r ≥ 0.85 were included. Expression patterns were further characterized using the Markov Cluster Algorithm (MCL) with an inflation value of 2 to identify clusters of genes with similar expression patterns.

### Behavioural analyses

Five cohorts of FIRE rats from different breeding pairs (Supplementary Table 1) underwent an 8-week behavioural testing pipeline (Figure 5a). From 6 weeks of age, rats were handled daily for 1-2 h a day to habituate them to the experimenter prior to testing, which started at 11 weeks. Starting at week 9, rats were placed on a food restricted diet (25 g/day) to maintain body weight at >85% of baseline throughout the study. All behavioural testing and data analysis were conducted blinded to genotype to minimise bias. Both male and female rats were included in all tasks to assess any potential sex differences. Full methodological details are provided in the Supplementary Information.

### Statistics

GraphPad Prism 10 software was used in statistical analyses. The critical significance level was set at α = 0.05, and *P* levels marked as \**P* < 0.05, \*\**P* < 0.01, \*\*\**P* < 0.001, \*\*\*\**P* < 0.0001. Details of statistical tests used and *P* values are located in the figure legends.

## Acknowledgements

We would like to express our gratitude to the dedicated staff at Bioresearch and Veterinary Services (BVS) at the University of Edinburgh for their invaluable support throughout our animal work. Their expertise and assistance played a crucial role in the success of this project. We gratefully acknowledge the support of the IRR core facilities at the University of Edinburgh. Flow cytometry data was generated with support from the IRR Flow Cytometry and cell sorting facility. Tissue processing and embedding were generated with support from the IRR Histology facility. We would also like to thank Josef Priller (University of Edinburgh) for providing the funding to purchase the gRNA-related reagents. This work was funded by the Simons Initiative for the Developing Brain.

## Author contributions

Experiments: E.K.G., X.L., Y.G., M.L., A.B., H.Z., C.P. Data analysis: E.K.G., X.L., D.DD., M.L., A.B., H.Z., O.T., C.P. Writing and editing of the manuscript: E.K.G., S.M.T., D.A.H., C.P. Project supervision and study conceptualisation: P.K., C.P.

