## Supplementary data for "Microglia are not required for normal postnatal brain development and cognitive function in the rat"

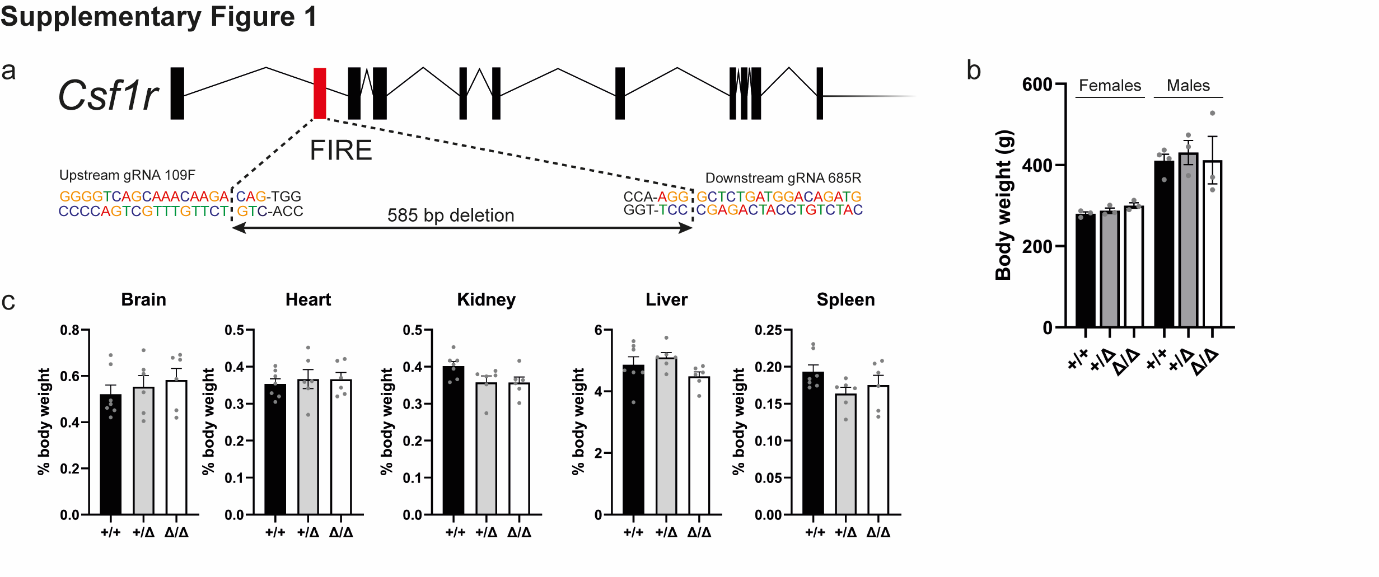

**Supplementary Figure 1**

**a)** Schematic showing the location of FIRE within rat *Csf1r* and gRNAs used to produce *Fireko* rats. Bases in black = protospacer adjacent motifs (PAM). **b)** Rats were weighed at 12 weeks of age. **c)** Organs were weighed from the same rats. Females: n = 3 per genotype. Males: n = 4 (+/+) and 3 (+/Δ, Δ/Δ). All graphs show mean + SEM. All *p* > 0.0724 via one-way ANOVA.

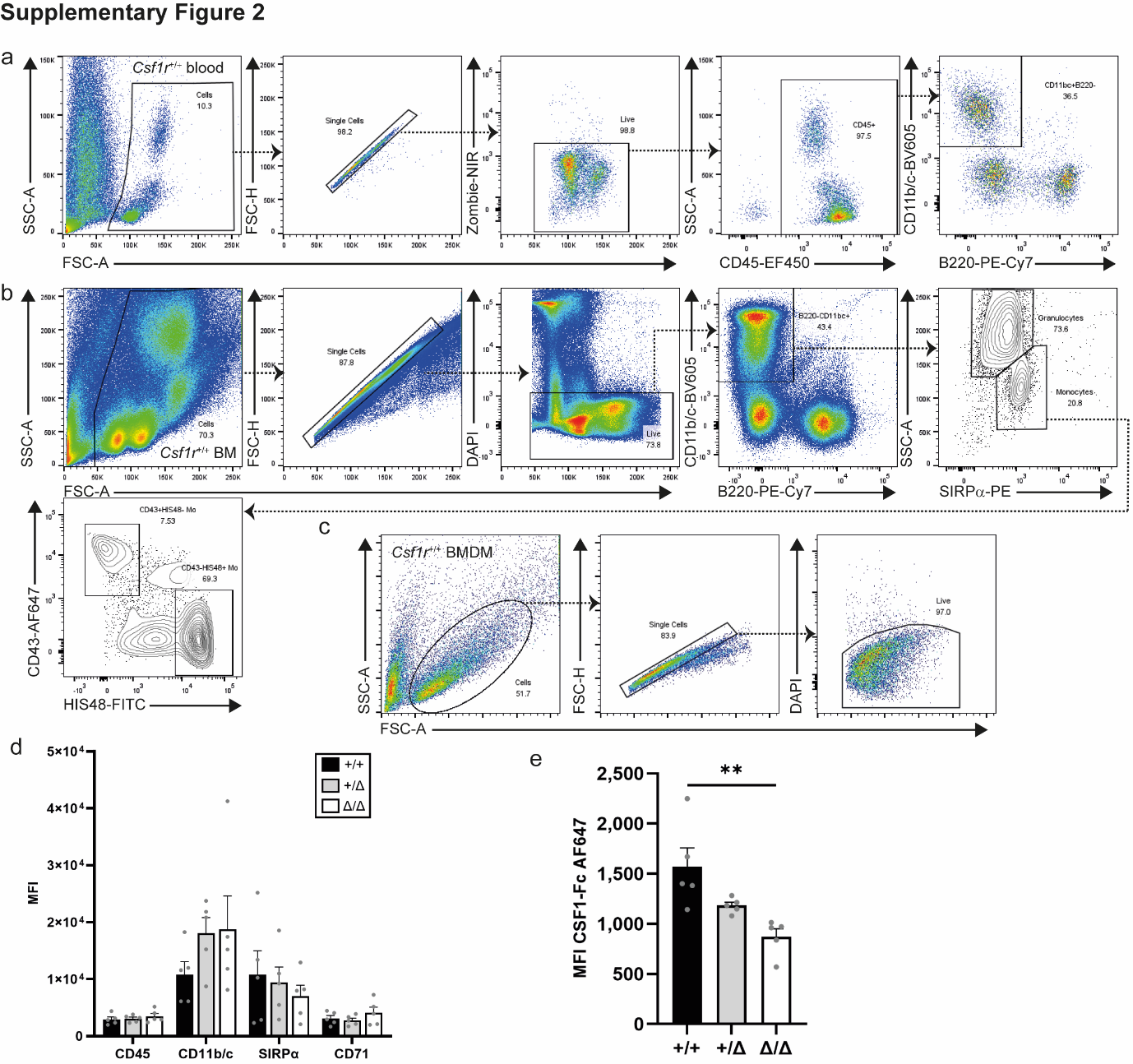

**Supplementary Figure 2**

**a)** Flow cytometry gating strategy for EDTA-blood (main article Figure 1a-b). **b)** Flow cytometry gating strategy for bone marrow (BM) in main article Figure 1c. **c)** Flow cytometry gating strategy for BM-derived macrophages (BMDM). **d)** Cryopreserved BM was placed in culture with 100 ng/mL recombinant human CSF1 and the BMDM analysed by flow cytometry after 7 days. MFI: mean fluorescence intensity. *n* = 5 rats per genotype. All *P* > 0.0767. **e)** BMDM were starved of CSF1 24 h before analysis by flow cytometry to examine CSF1-Fc binding as a marker of CSF1R expression. *n* = 5 rats per genotype. *P* = 0.0039 (**). All graphs show mean + SEM and *P* values were determined by one-way ANOVA with Tukey’s multiple comparisons test. Each data point represents an individual animal.

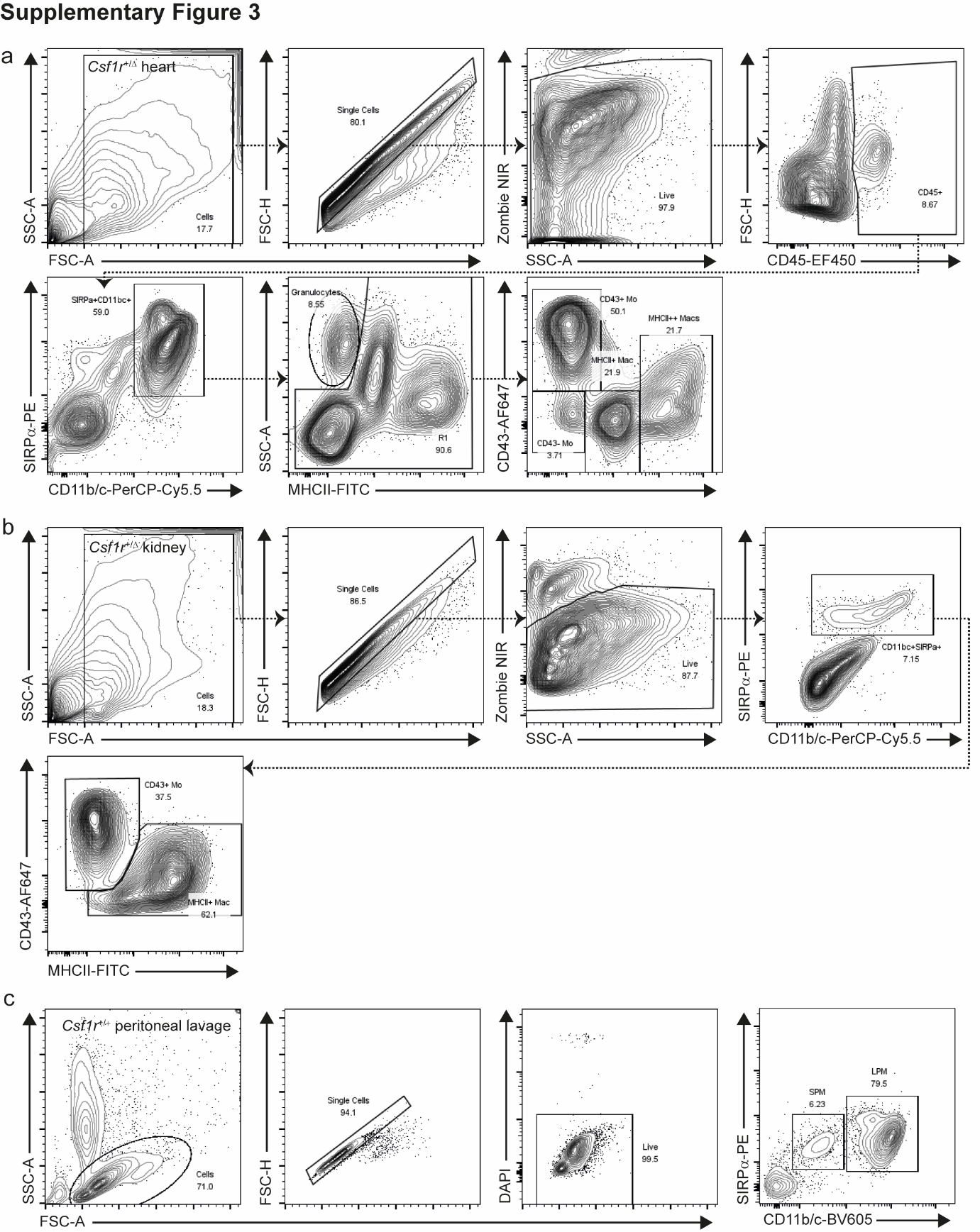

**Supplementary Figure 3**

Flow cytometry gating strategies for **a)** hearts (main Figure 2a), **b)** kidneys (main Figure 2b) and **c)** peritoneal lavage (main Figure 2c-d). Mo: monocyte, Mac: macrophage, SPM: small peritoneal macrophage, LPM: large peritoneal macrophage.

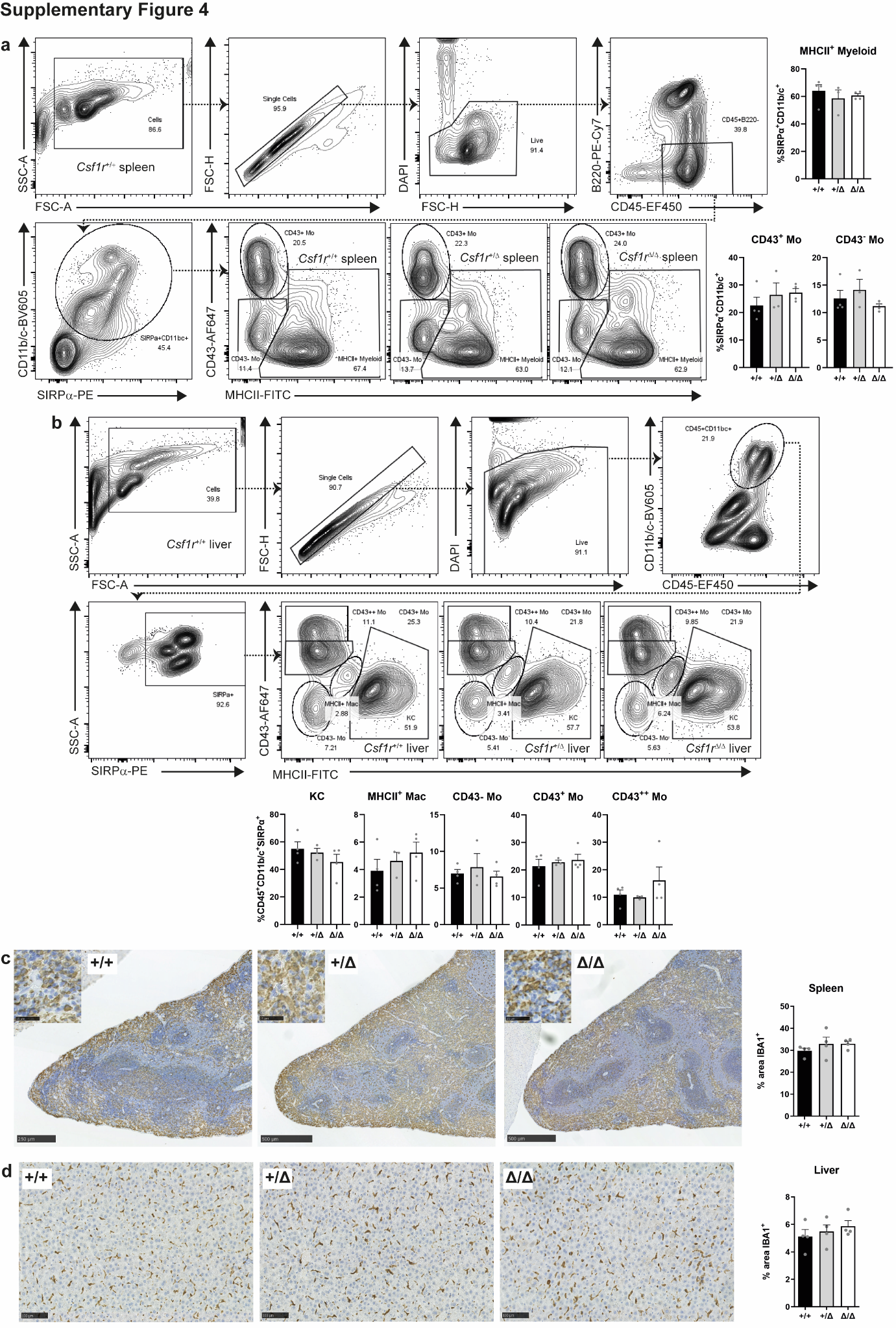

**Supplementary Figure 4**

Male and female rats aged 8-12 weeks were culled via a rising concentration of CO_2_ and single cell suspensions of enzymatically digested spleens and livers were analysed by flow cytometry. *n* = 3-4 rats per genotype from 3 independent experiments. Isotype and fluorescent minus one controls were used to determine gates. **a)** Gating strategy and representative flow cytometry profiles of spleens and **b)** livers. Mo: monocyte, KC: Kupffer cell, Mac: macrophage. Formalin-fixed paraffin-embedded organs from a different cohort of rats were stained with an antibody against IBA1. *n* = 4 rats per genotype **c)** Representative IBA1 staining of spleens. Scale bar = 500 µm and 25 µm (inset). **d)** Representative IBA1 staining of livers. Scale bar = 100 µm. All graphs show mean + SEM. All *P* > 0.3271 via one-way ANOVA.

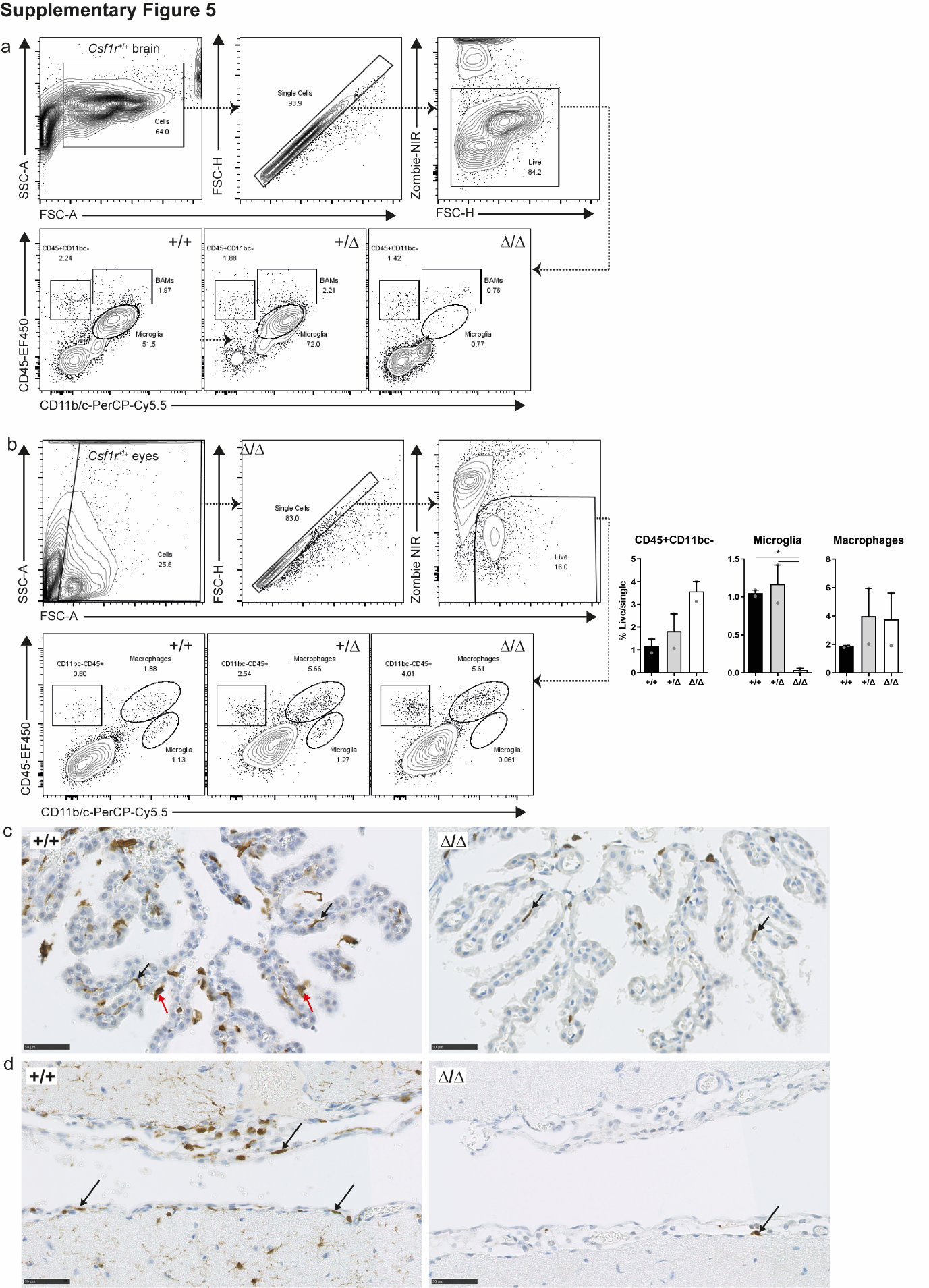

**Supplementary Figure 5**

**a)** Flow cytometry gating strategies for brains (main Figure 3c). **b)** Eyes from rats aged 2-3 months old were digested with an enzyme cocktail and analysed by flow cytometry. Graphs show mean + SEM. *P* = 0.0332 (* +/+ vs. Δ/Δ) and 0.0245 (* +/ Δ vs. Δ/Δ) via one-way ANOVA with Tukey’s multiple comparisons test. Formalin-fixed paraffin-embedded brains from male and female rats aged 2-3 months were stained with an antibody against IBA1. **c)** Black arrows: choroid plexus stromal macrophages. Red arrows: choroid plexus epiplexus macrophages. **d)** Black arrows: meningeal macrophages from the cortex. Two adjacent brain sections from each paraffin block are shown. Images are representative of 5 +/+ and 6 Δ/Δ.

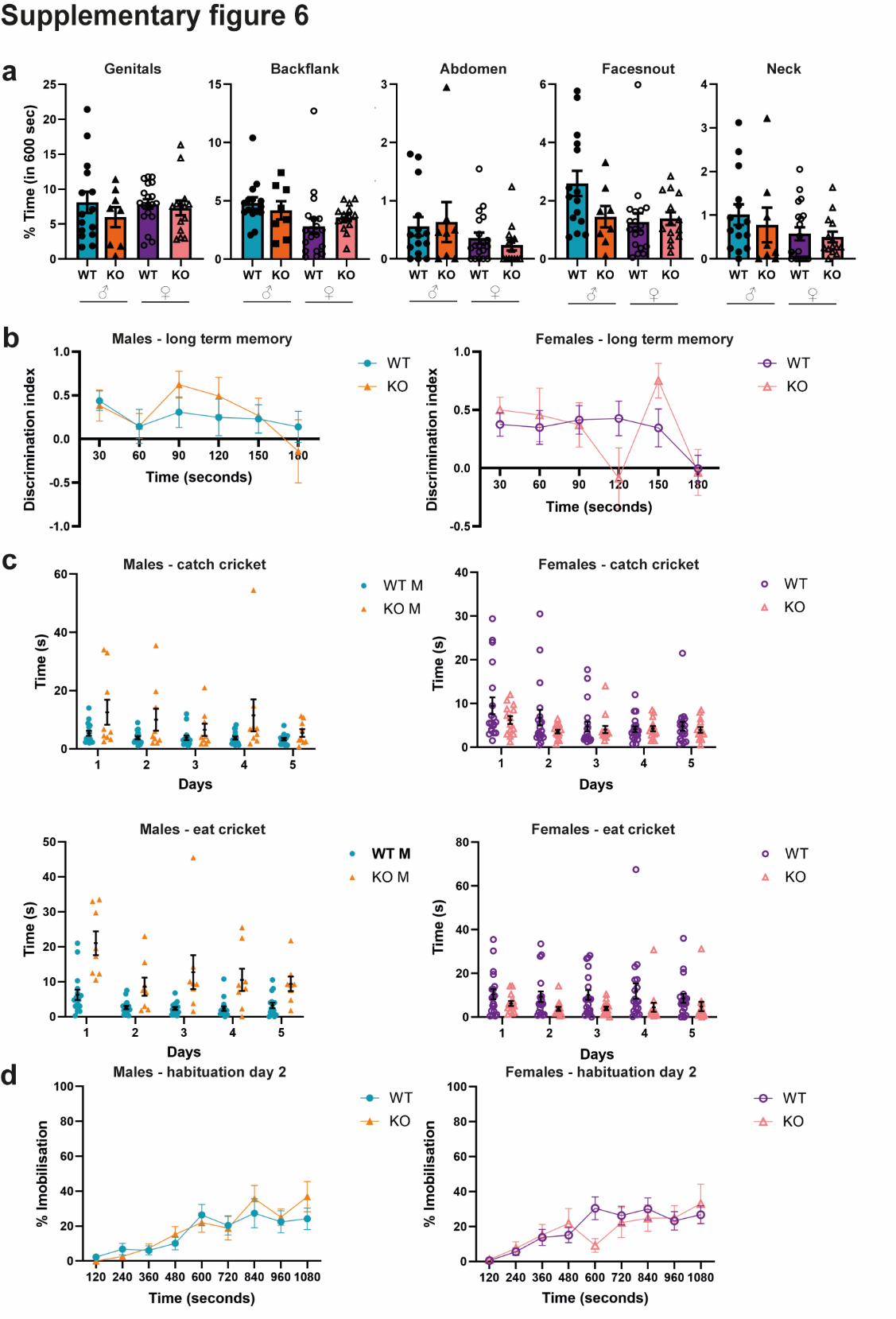

**Supplementary Figure 6**

**a)** Regions of the body sniffed in the one trial social task (main Figure 5) **b)** Long term memory object recognition of male and female 12 week old rats. **c)** Latency to catch and eat a live cricket in prey capture assessment of 16 week old rats. *n* = 9-19 rats per genotype and sex. **d)** Day 2 of fear conditioning assessment of 18 week old rats. *n* = 9-19 rats per genotype and sex. Graphs show mean ± SEM and all *P* values > 0.05 via two-way ANOVA with Tukey's multiple comparisons test (except for eat cricket which was analysed by mixed-effect model as 2 rats did not eat a cricket every day). Sexes were separated for illustrative purposes. WT: *Csf1r*^+/+^/ *Csf1r*^+/ΔFIRE^, KO: *Fireko*.

**Supplementary Table 1 -** Rat cohort numbers and genotypes used for behaviour

pipeline.

| **Cohort** | **Genotype and sex** |
| --- | --- |
| 1 | +/+ = 4 females  +/Δ = 4 females  Δ/Δ = 4 females |
| 2 | +/+ = 3 males  +/Δ = 3 males  Δ/Δ = 3 males |
| 3 | +/+ = 2 males & 2 females  +/Δ = 2 males & 2 females  Δ/Δ = 2 male & 1 female |
| 4* | +/+ = 3 females  +/Δ = 4 males & 2 females  Δ/Δ = 2 males & 2 females |
| 5 | +/+ = 2 males & 2 females  Δ/Δ = 2 males & 8 females |
| Total males | +/+ = 7  +/Δ = 9 (16^a^ control)  Δ/Δ = 9^b^ |
| Total females | +/+ = 11  +/Δ = 8 (19 control)  Δ/Δ = 15^ab^ |

* cohort did not undergo auditory fear conditioning

^a^ one rat did not complete social task due to aggression

^b^ one rat did not eat the cricket

**Supplementary methods**

*One trial social task*

Test rats were randomly assigned a sex matched wild type stimulus rat of a similar age for the trial. Each stimulus rat was used only once. All rats, both test and stimulus, were habituated to a 1 m x 1 m arena with white walls and a layer of sawdust covering the floor. Light levels were maintained at ~20 lux using a standing lamp. All habituation and trial sessions were video recorded. On day 1 of habituation, each cage group of rats was placed in the arena together for 10 min. On day 2, each rat was individually habituated to the arena for 10 min, in the reverse order of previous day’s cage group sessions. This allowed all rats to become familiar with each other’s scent in the arena. Faeces were removed between sessions, but the same sawdust was used for all rats. On the trial day, the test rat was placed into the arena for 5 min before the assigned stimulus rat was introduced into the corner opposite to where the test rat had moved. The two rats were then allowed to interact for 10 min. If any fighting or wrestling occurred, the rats were separated. At the end of the session, the test rat was always removed from the arena first.

For analysis, the occurrence and duration of different behaviours were coded manually from video recordings using Behavioural Observation Research Interactive Software (BORIS; ^1^.

*Object recognition*

Rats were habituated to a 1 m x 1 m arena with black panelled walls, a lino floor with Velcro at two points and no specific cues. Light levels were maintained at 10-15 lux using a standing lamp. All sessions were video recorded. For the short-term memory object recognition (STM OR) task, Day 1 of habituation involved each cage group spending 10 min in the arena. On Day 2 and 3, each rat was placed individually in the arena for 10 min, followed by 5 min in a blackout holding bucket. On Day 4, each rat was exposed to 2 identical objects in the arena for 10 min, followed by 5 min in the holding bucket. One of the objects was then replaced with a novel object, and the rat was returned to the arena for a 3 min test session. The objects were chosen to be visually distinct and engaging for the rats; in STM OR these included a ceramic teddy bear and a mirror weighted ‘XOXO’ sign.

The long-term memory object recognition (LTM OR) task followed the same 3 day habituation protocol. On Day 4, each rat was exposed to 2 identical objects for 10 min, followed by 5 min in the holding bucket. 24 h later, the rats were returned to the arena for a 3 min test with one object changed. LTM OR objects included a tall pink lemon juicer and short pink heart shaped tea light holder. Between rats, the arena and objects were cleaned with 70% ethanol to remove scent cues. Object interactions were manually coded in 30 s intervals over the 3 min test using BORIS^1^.

*Prey Capture*

Protocol was adapted from the ‘Hunting Test’ in ^2^. Rats were habituated in a 1.5 m x 1.5 m arena with white walls and a plain white floor. Crickets (large adult silent crickets, Livefoods Direct Ltd) were kept in a separate room to the testing area. All sessions were video recorded. On Day 1 of habituation, rats explored the arena for 30 min with their cage mates. On Day 2, each rat was individually placed in the arena for 10 min. On Day 3, each rat was given a dead cricket in their home cage to acclimate to the taste. Crickets were killed by applying pressure to the head. During this session, each rat was placed into a holding cylinder in one corner of the arena whilst a dead cricket was positioned in the opposite corner. After 15 sec, the rat was released and allowed to explore the arena for 10 min. The time taken to eat the cricket was recorded. On Days 4 and 5, this procedure was repeated with a new dead cricket. After each session, the rat was placed in a blackout holding bucket for 2 min.

Trial Days 1, 2, 3 and 4 involved placing the rat in the holding cylinder for 15 s, with a live cricket placed in the opposite corner. The rat was then released and allowed up to 2 min to hunt the cricket. This was repeated 4 times per day, using each corner of the arena as the starting point. Between each of the 4 trials, the rat was placed in the blackout holding bucket. Trial day 5 followed the same procedure, but the live cricket was placed in a pseudorandom location. For each trial, 3 measurements were recorded: the time taken to intercept, catch and eat the cricket. Between trials, the arena was cleaned with water to remove any faces or urine and minimise distraction.

*Auditory fear conditioning*

Two distinct contexts located inside sound-isolation boxes (Coulbourn Instruments, Whitehall, PA, USA) were used, with different appearances and odours. The testing context had black and white striped curved walls, a house light (5 lux), a flat floor and mint odour. The main room lights were fully on. This context was cleaned with 70% ethanol between each trial. The conditioning context had plain walls, two house lights, no odours and a grid shock floor. The main room lights were off with a standing lamp providing lower light levels. This arena was cleaned with Azowipes (Synergy Healthcare) to prevent lingering scents. A video camera mounted above each context recorded the sessions.

Context habituation involved each rat exploring the testing context for 20 min on 2 consecutive days. Before returning to the home-cage, each rat was isolated in a holding cage for 5 min, to avoid any social transmission of fear. On Day 3, rats were placed in the conditioning context where they experienced 3 pairings of a conditioned stimulus (CS; 30s 5 Khz tone at 75 dB) co-terminating with a 1 sec scrambled foot shock unconditioned stimulus (US; 0.7 mA). The CS-US pairings were presented 2 min after entering the context and were separated by a
60 sec inter-tone interval (ITI). Recall and extinction of the conditioned response were tested on Days 4 and 5. Each rat was given 2 min to explore the testing context, then presented with 13 CS with a 30 sec ITI. Percent time spent freezing was quantified using FreezeFrame 5 (ActiMetrics).
